# Swimming speed of schooling fish controls social interaction strength in open-loop immersive virtual reality

**DOI:** 10.64898/2026.01.08.698350

**Authors:** Ramón Escobedo, Justine Reynaud, Stéphane Sanchez, Clément Sire, Guy Theraulaz

**Affiliations:** Centre de Recherches sur la Cognition Animale, Centre de Biologie Intégrative, CNRS, Université de Toulouse – Paul Sabatier, Toulouse, France; Institut de Recherche en Informatique de Toulouse (IRIT), Université Toulouse Capitole, Toulouse, France; Laboratoire de Physique Théorique, CNRS, Université de Toulouse – Paul Sabatier, Toulouse, France; Departamento de Matemáticas, Universidad Carlos III de Madrid, Leganés, Madrid, Spain

## Abstract

Collective motion in fish schools emerges from local attraction, alignment, and repulsion rules whose strengths vary with behavioural and biomechanical context. Although swimming speed is known to shape turning capacity, spatial positioning, and information flow, its causal influence on social interaction strength remains poorly quantified. Using an open-loop immersive virtual-reality system, we expose freely swimming *Hemigrammus rhodostomus* to a virtual conspecific moving along circular trajectories at six controlled speeds. This approach isolates the effect of neighbour speed on attraction, alignment, vertical positioning, and wall avoidance. Across all speeds, real fish reliably track the virtual fish and adjust their own speed toward that of the partner. Behavioural distributions reveal systematic changes in spatial positioning and perception angles, with fish shifting from leader-like to follower-like configurations as virtual-fish speed increases. A data-driven 3D self-propelled particle model, in which only the amplitudes of interaction functions are modulated, quantitatively reproduces these patterns. Model-based reconstructions shows that higher partner speed weakens both horizontal and vertical attraction, strengthens alignment, and increases wall repulsion. These results demonstrate that fish tune their social-interaction rules in a speed-dependent manner, revealing swimming speed as a key control parameter shaping coordination in minimal groups and, potentially, in larger schools.

## 1 Introduction

Collective motion in fish schools emerges from simple local interactions yet displays remarkable coordination, suggesting that individuals continuously adjust attraction, repulsion, and alignment responses in relation to their neighbours [1–5]. While many classical models assume constant swimming speed for simplicity, a growing body of empirical and theoretical work shows that individual speed is highly variable and profoundly shapes local and global collective dynamics [3, 6–8]. Experimental observations in diverse fish species and biomimetic robots demonstrate that instantaneous speed fluctuates within and across individuals and that these fluctuations influence turning capacity, alignment strength, and local information flow [2, 5, 8–10]. Specifically, turning dynamics depend on speed through fundamental kinematic constraints: at higher speeds, individuals turn more slowly, which couples speed to responsiveness and imposes structure on interaction patterns [2, 3, 5, 8, 11–14]. This link between speed, turning, and social responsiveness has major consequences for emergent group organisation. Agent-based models incorporating variable speed reveal that speed variability alone can enhance or destroy polarization, shift the location of the order-disorder transition, and induce bistable collective states that do not occur in fixed-speed models [14]. These findings highlight that speed is not a secondary attribute of motion but a central behavioural parameter governing the emergent collective patterns.

Recent empirical work further shows that social interactions themselves depend on the relative speeds of neighbours. Using force-map analyses of schooling fish, it has been demonstrated that individuals tend to align only with neighbours that swim faster than themselves, while slower neighbours are effectively ignored or may even produce apparent anti-alignment signals [15]. This selective attention to faster neighbours creates transient, speed-based leader-follower relationships that continuously reorganize the flow of social influence within the group [5]. These interactions amplify local speed differences, enabling faster individuals to transiently lead collective movement, while slower individuals fall behind and have reduced impact on group orientation. Similar conclusions have been drawn from experiments employing highly social robotic fish, which revealed that individual speed strongly structures spatial positioning and group-level patterns, with faster agents tending to occupy frontal positions and exert disproportionate social influence [8]. Together, these studies show that neighbour speed determines not only the magnitude but also the direction and selectivity of social responses.

Speed thus acts as a behavioural signal that modulates the strength of social interactions, influences leadership, and shapes the geometry and cohesion of fish schools [3, 8, 13]. At the group level, variations in speed affect polarization, spatial extent, and stability of collective states, while at the individual level, they modulate alignment, attraction, and the distribution of social attention [6, 9, 10, 16, 17]. Movement-ecology studies across taxa show that groups with higher mean speed tend to be more polarized and cohesive, whereas groups with greater speed variability exhibit lower polarization and increased susceptibility to disorder [2–4, 6–8, 16, 17]. This sensitivity of collective behaviour to speed and speed variability emphasizes the need for controlled experiments capable of isolating the causal impact of speed on social responses.

In this context, the present study experimentally examines whether and how the swimming speed of a neighbour shapes the social interactions of a focal fish. Using an open-loop immersive virtual reality system, we exposed freely swimming *Hemigrammus rhodostomus* to a virtual conspecific moving along identical circular trajectories at different constant speeds. This approach makes it possible to manipulate neighbour speed independently of the real fish’s behaviour, thereby isolating its causal effect on alignment, attraction, and spatial positioning [18]. By analysing the responses of real fish across a systematically varied range of virtual speeds, we test whether interaction strength increases with neighbour speed, whether spatial positioning changes accordingly, and whether speed alone can reorganize leader-follower dynamics in minimal two-agent groups. To complement these experiments, we used a data-driven 3D self-propelled particle model to estimate how the strengths of social interactions vary with speed and to assess whether speed-dependent modulation of interaction rules is sufficient to reproduce the behavioural patterns observed in the virtual reality experiments. Situating our approach within the growing literature on speed-mediated interactions demonstrates that speed is not merely a kinematic output but a key driver of social responsiveness, selective attention, and emergent coordination in animal collectives [19, 20].

## 2 Methods

### 2.1 Laboratory experiments

#### 2.1.1 Study species

Rummy-nose tetras (*Hemigrammus rhodostomus*) were purchased from Amazonie Labège in Toulouse, France. Fish were kept in 16 L aquariums on a 12:12 hour, dark:light photoperiod, at 24.9° C (± 0.8° C) and were fed *ad libitum* with fish flakes. The average body length of the fish used in these experiments is 3.1 cm.

#### 2.1.2 Virtual reality platform

We conducted our experiments using a custom virtual reality platform [18], designed for studying real-time interactions between freely swimming fish and virtual conspecifics in open- and closed-loop configurations.

The core of the setup is a hemispherical acrylic bowl (diameter 52.4 cm) containing 15 L of water (depth 14.73 cm at the centre) and positioned directly over a DLP LED projector, which was responsible for displaying the anamorphically rendered virtual fish on the bowl’s wall (Fig. 1). To track fish movements, an Intel RealSense D435 depth camera was mounted 47.9 cm above the water surface. Robust tracking under controlled lighting was achieved using infrared illumination (8 IR lamps, 850 nm). A dedicated workstation (Dell Precision 3640 with NVIDIA RTX 3070) performed all real-time processing (3D tracking, simulation of virtual fish trajectories, and rendering).

**Figure 1:**
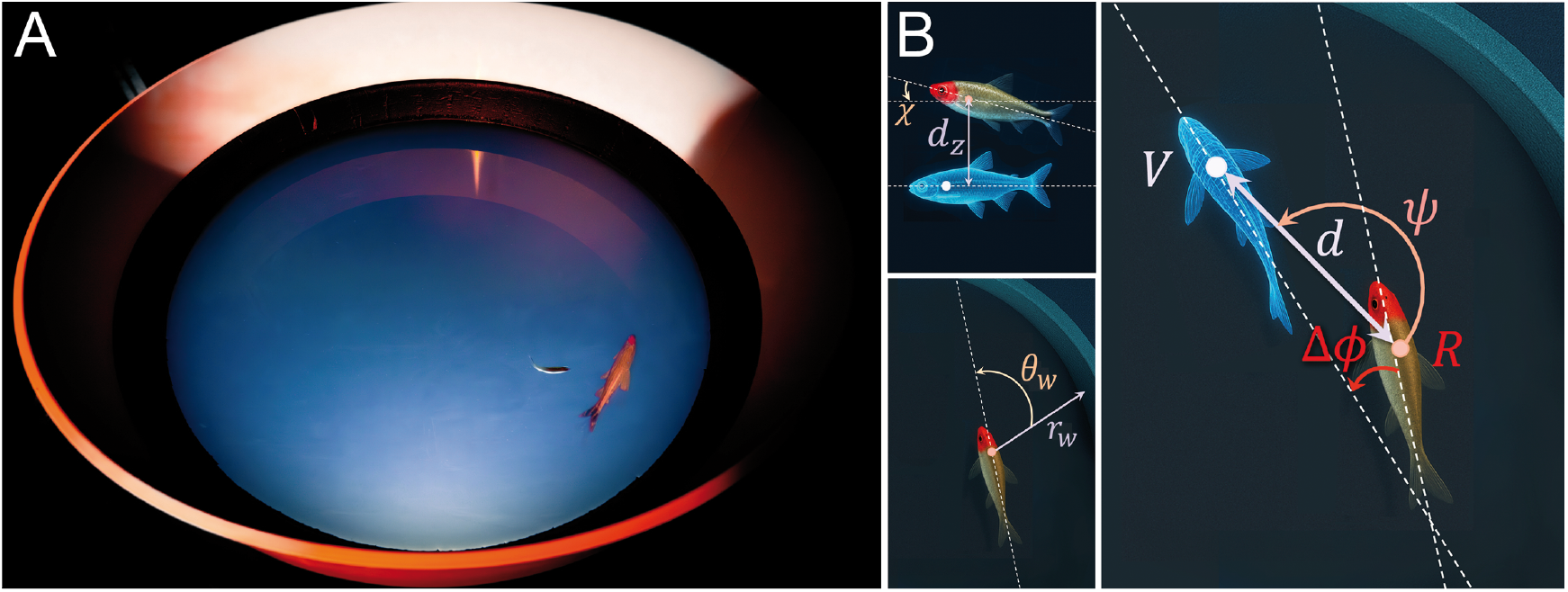
Virtual-reality setup and geometric variables describing real-virtual fish interactions. (A) Virtual reality bowl with a real fish (small) and the 3D anamorphic image of a virtual fish (large) projected on the bowl’s wall. (B) Variables describing the distance *r*_*w*_ and relative orientation of the real fish to the wall *θ*_*w*_, the horizontal and vertical distances between individuals *d* and *d*_*z*_, the viewing angle *ψ*, the heading angle *ϕ*, the relative orientation Δ*ϕ*, and the elevation angle *χ*.

The system’s architecture comprised three main software components: a 3D tracking pipeline written in Python, a trajectory simulator implemented in C++, and a Unity-based rendering engine. For this investigation, the system operated in open-loop mode, whereby the virtual fish’s position and heading were fixed to circular trajectories at constant speed, constant distance to the border, and constant depth, independently of the real fish’s instantaneous position and speed. Communication of virtual fish positions to the real one was handled via UDP at a frequency of 30–90 Hz, guaranteeing precise and biologically meaningful interactions. This identical configuration was used to study the behaviour of a single real fish reacting to a virtual one swimming at different constant speeds, maintaining the same structured background and lighting conditions in all speed conditions.

#### 2.1.3 Experimental procedure

We conducted open-loop experiments to measure the effect of swimming speed on social interactions in fish, by studying the behaviour of a single real fish in the presence of a virtual conspecific that swims at different constant speeds. The virtual fish moved along circular trajectories centred in the tank, with a radius of 15 cm (corresponding to a distance of 5.4 cm to the tank wall) and at a depth of 5 cm, close to the average swimming depth observed for pairs of real fish [21]. Similar circular motion patterns were previously observed in experiments with real fish swimming alone or in pairs in the same experimental setup.

Six different constant swimming speeds of the virtual fish were tested, ranging from v_1_ = 5 cm/s (the typical speed of two real fish swimming together) to v_6_ = 15 cm/s (considered high for *H. rhodostomus* when swimming in circular tanks with 7 cm of water depth). Intermediate speeds were 7.5, 10, 11.67, and 13.33 cm/s. For each virtual fish speed condition, 7 to 9 one-hour experimental replicates were performed.

#### 2.1.4 Geometric and kinematic variables used to quantify real-virtual fish interactions

Because fish rarely change their swimming depth and remain largely confined to a horizontal plane near the surface, we simplify the notation as follows. Three-dimensional quantities are indicated with a superscript 3D, horizontal (2D) quantities are written without any superscript, and vertical components are denoted with a subscript *z*.

The experimental bowl is the lower cap of a perfect sphere of radius *R*_0_ = 26.19 cm, filled with 15 litres of water, resulting in a central depth of *h*_water_ = 14.73 cm. The origin of coordinates is placed at the water surface on the bowl’s vertical axis. This axis defines the *z*-axis, oriented downward so that depth values are always positive and increase with distance from the surface. At depth *z*, the radius of the horizontal cross-section is such that 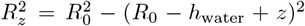, i.e., *R*_*z*_ = 0 at the bottom of the bowl, where *z* = *h*_water_.

The fish position and velocity are given by the vectors 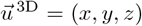 and 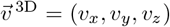. We define the distance to the bowl vertical axis as *r* = ∥(*x, y*)∥ and the distance to the horizontal boundary at depth *z* as *r*_w_ = *R*_*z*_ − *r*. For consistency with previous work, we refer to this horizontal boundary as the *wall*.

The horizontal velocity is 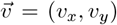, and the full 3D-speed and its horizontal component are 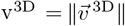 and 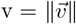, respectively. The fish heading in the horizontal plane is defined as *ϕ* = atan2(*v*_*y*_, *v*_*x*_) ∈ (−*π, π*], while the elevation angle is *χ* = atan(*v*_*z*_*/*v) ∈ (− *π/*2, *π/*2). The heading is used to compute the angle of incidence to the wall, *θ*_w_ = *ϕ* − *θ*, where *θ* = atan2(*y, x*) is the positional azimuth. By convention, positive headings are anticlockwise, and positive (negative) elevation indicates upward (downward) motion.

When swimming with the virtual conspecific, the relative state of a fish is determined by the distance to the other fish, the angle with which it perceives the other, and the difference of alignment. For a pair of fish (*i, j*), the 3D distance is 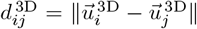, the horizontal distance *d*_*ij*_ = ∥(*x*_*i*_, *y*_*i*_) − (*x*_*j*_, *y*_*j*_)∥, and the vertical distance 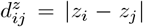. The viewing angle is the angle that the fish *i* has to turn to point towards fish *j*, given by *ψ*_*ij*_ = atan2(*y*_*j*_ − *y*_*i*_, *x*_*j*_ − *x*_*i*_) − *ϕ*_*i*_. Note that, in general, *ψ*_*ij*_ ≠ *ψ*_*ji*_. Finally, alignment differences in the horizontal and vertical planes are respectively given by *ϕ*_*ij*_ = *ϕ*_*j*_ − *ϕ*_*i*_ = − *ϕ*_*ji*_ and *χ*_*ij*_ = *χ*_*j*_ − *χ*_*i*_ = −*χ*_*ji*_.

We define the *geometrical leader* as the individual who has to turn a larger absolute angle |*ψ*_*ij*_| to face its partner. The other fish is then the *geometrical follower*. As only dyads are considered here, indices are omitted when no ambiguity arises, writing *d, ψ*, Δ*ϕ*, Δ*χ* instead of *d*_*ij*_, *ψ*_*ij*_, *ϕ*_*ij*_, *χ*_*ij*_ (see Fig. 1).

### 2.2 3D model of fish swimming and interaction rules

We developed a three-dimensional model to quantitatively compare with experiments and explain the observed collective behaviours. This model extends the successful 2D framework originally established by Calovi et al. [22], which has proven effective in describing social interactions in confined geometries [23–25]. Adapted here to pairs of fish swimming in a spherical bowl, the model describes the continuous motion of a fish in terms of its behavioural state, which is defined by a 9-tuple of kinematic and social variables (v, v_*z*_, *r*_w_, *θ*_w_, *z, d, d*_*z*_, *ψ*, Δ*ϕ*).

The dynamics of these variables combine biomechanical and environmental constraints with social interactions with the partner. The equations are grounded in principles of self-organization, symmetry, and collision avoidance, with their specific functional forms quantitatively inferred from experimental data using a reconstruction procedure adapted from flat-arena studies to our curved 3D geometry [22, 26]. Details of the specific 3D procedure are given in [21].

The model decomposes fish motion into horizontal and vertical dynamics:

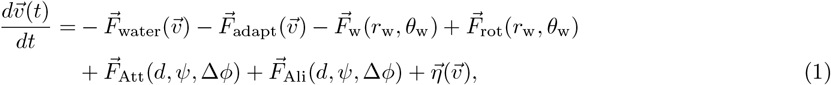

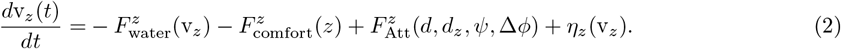

In the horizontal plane, the instantaneous acceleration 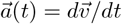 results from six components: (**i**) an effective friction emerging from hydrodynamic effects and the tendency of fish to swim at a preferred speed 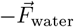, (**ii**) a tendency of a fish to adapt its speed to that of its neighbour (or to the average speed 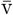 of its neighbours, for more than one neighbour), 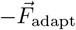, (**iii**) a radial centripetal repulsion from the wall of the bowl 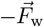, (**iv**) a rotational force of deviation induced by the wall 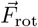, and the social forces of (**v**) attraction 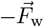, and (**vi**) alignment 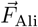. The vertical acceleration 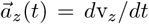 depends on three components: (**i**) the friction acting on the vertical speed 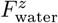, resulting in the fish tending to swim horizontally, (**ii**) a tendency to swim at a preferred depth, resulting from the interaction with the water surface and the bowl 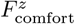, and (**iii**) the social vertical attraction towards a partner’s depth 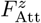. Additionally, a noise term is added to each component, 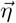 and *η*_*z*_ respectively, corresponding to the fish’s spontaneous variability of individual behaviour.

The horizontal velocity can be decomposed into the parallel and perpendicular components of its instantaneous heading, given by the 2D velocity vector. Defining the unit vectors 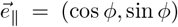 and 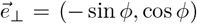, the velocity in the horizontal plane is given by 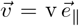. The horizontal acceleration is written as 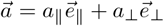, where *a*_∥_ and *a*_⊥_ are the longitudinal (speed-changing) and lateral (turning) acceleration components, respectively.

The scalar components of the acceleration are then as follows:

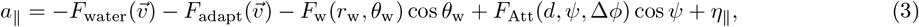

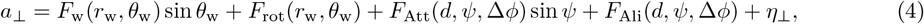

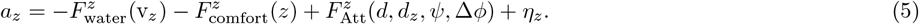

This kinematic decomposition allows us to incorporate the fundamental symmetry constraints and preserve the directional reactions of fish (right-left symmetry), which was shown to consistently hold for the considered fish species [22]. The interaction functions can be extracted from the two real-fish experiments through an optimization procedure that minimizes the error between the experimental measures and the model predictions [21, 22, 26]. Additional details completing the formulation of the model are provided in the electronic supplementary material, including the analytical expressions of the interaction functions.

Then, for each speed condition, the strength of the social interactions and the effect of the bowl’s wall are adjusted to reproduce the behaviour of the real fish. The adaptation is minimal and consists of applying a specific multiplicative factor to each of the four interaction functions *F*_Att_, 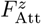, *F*_Ali_, and *F*_w_. Simulation results obtained with these adjusted functions for speeds v_1_ to v_6_ are presented alongside the corresponding experimental data.

## 3 Results

Figure 2 shows six short trajectory excerpts of both the real and the virtual fish for the different values of the virtual fish’s speed.

**Figure 2:**
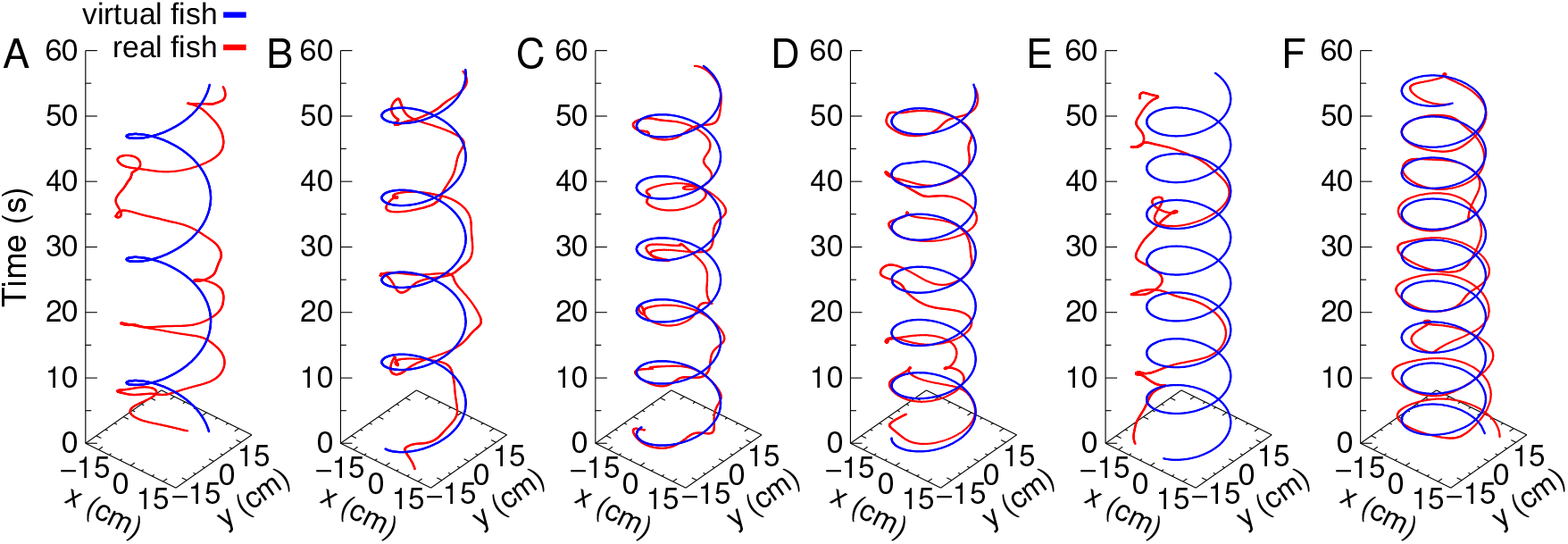
Real and virtual fish trajectories across six swimming speed. Trajectories of a real fish (red) interacting with a virtual fish (blue) that swims along a circular trajectory at constant speed, fixed distance from the tank wall (5.4 cm), and constant depth (5 cm). Panels show six swimming speeds of the virtual fish, from v_1_ to v_6_: (A) 5, (B) 7.5, (C) 10, (D) 11.67, (E) 13.33, and (F) 15 cm/s.

In all cases, the real fish follows the virtual fish’s trajectory quite closely. At lower speeds (Fig. 2A, B, C, electronic supplementary material, movie S1), it is frequent that the real fish deviates from the virtual fish’s path, forming loops and more sinuous trajectories, a behaviour that can also be observed at higher speeds (Fig. 2E, electronic supplementary material, movie S2). When swimming with the virtual fish, the real one can be either on the inner side of the circular path (as in loops 3, 7–9 in Fig. 2F), where the image representing the virtual fish is projected on the same side of the bowl as its presumed location with respect to the real fish, or on the outer side (as in loops 1, 2, and 4–6), where the image appears on the opposite side of the bowl, so that the virtual fish is seemingly positioned between the projection wall and the real fish.

### 3.1 Real fish adjust their swimming behaviour to virtual conspecific speed

Figure 3 shows the mean values of four key observables characterizing the behaviour of the real fish: its speed v, its horizontal distance to the wall *r*_w_, its swimming depth *z*, and its horizontal distance to the virtual fish *d*. The corresponding regression analyses are shown in Table S1.

**Figure 3:**
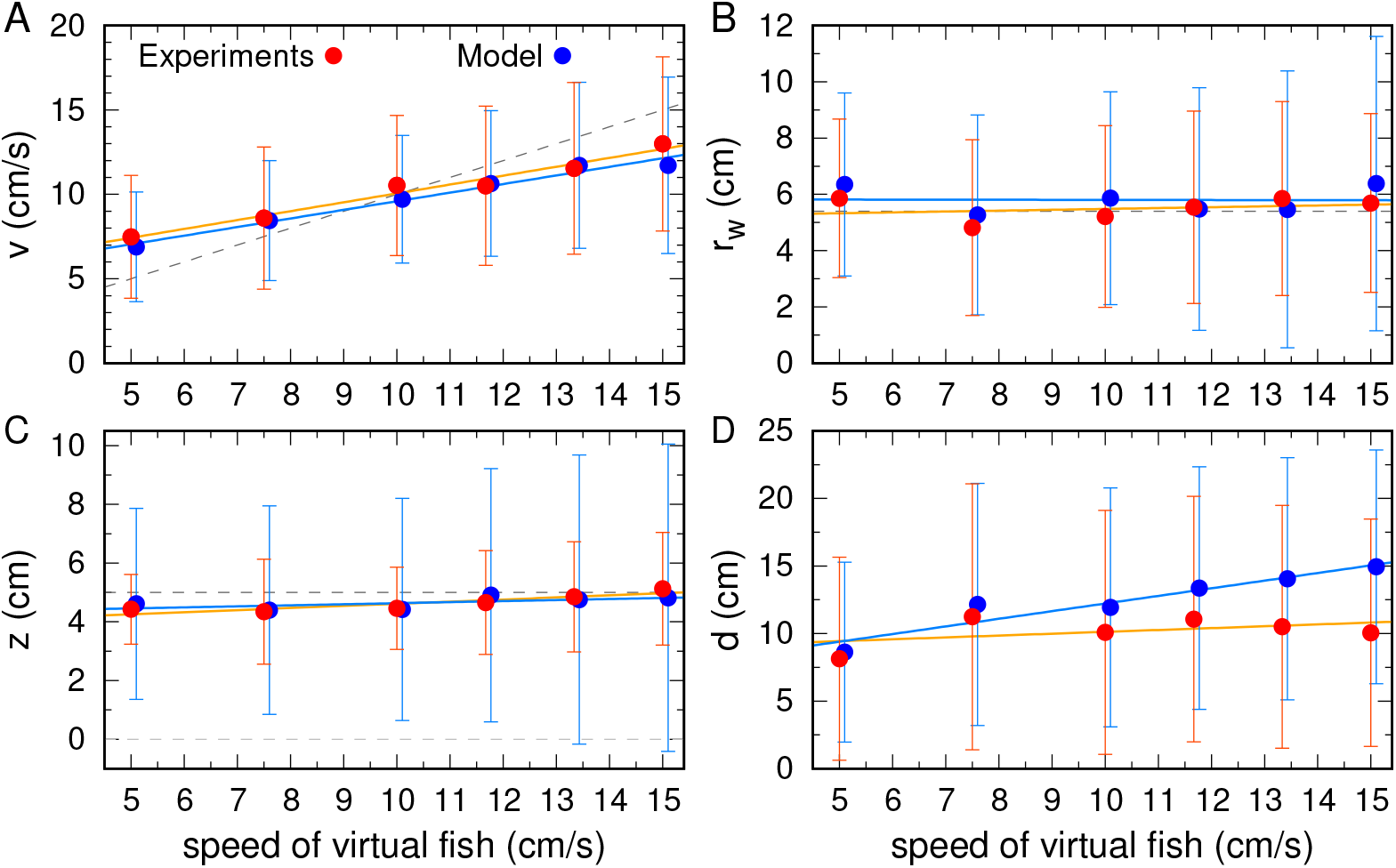
Behavioural responses of real fish to virtual-fish speed. Mean behavioural metrics of the real fish as a function of the virtual fish’s swimming speed (v_1_ to v_6_: 5, 7.5, 10, 11.67, 13.33 and 15 cm/s). (A) Swimming speed v, (B) distance to the wall *r*_w_, (C) distance between individuals *d*, and (D) swimming depth *z*. Red symbols correspond to experimental data, and blue symbols to numerical simulations. Mean values are shown as circles; vertical bars indicate the standard deviation of the corresponding variable. Dashed grey lines in panels A–C represent the constant values of *v, r*_w_, and *z* of the virtual fish. For clarity, circles and bars for the numerical simulations have been shifted by 0.1 cm/s to the right.

The first observation is that the mean speed of the real fish shows a clear quasi-linear increase with that of the virtual partner (Fig. 3A), indicating a tendency to adopt the partner’s speed. In addition, the data reveal a critical virtual-fish speed, slightly above v_3_ = 10 cm/s, below which the real fish is faster than the virtual one, and above which the real fish is slower than the virtual one (see the dashed gray line in Fig. 3A corresponding to *y* = *x*). Such a pattern is consistent with the existence of an intrinsic preferred speed of comfort close to this critical value.

The regression line of the mean distance to the wall practically overlaps the constant value imposed on the virtual fish (Fig. 3B). For depth, the regression line is also almost horizontal, showing that the real fish is on average only slightly below the virtual one (Fig. 3C). Finally, the regression line of the mean distance is also practically horizontal, except in the slowest case v_1_ = 5 cm/s, where the real fish is on average slightly closer than at the other speeds (Fig. 3D).

The mean behaviours observed in Fig. 3 result from underlying probability density functions (PDFs) that are shaped by the speed of the virtual fish. Figure S1 in the electronic supplementary material presents the PDFs of individual behavioural variables of the real fish, comparing the experimental results (left column, in red) to the model simulations (right column, in blue).

The PDF of the speed (Fig. S1A) broadens and shifts toward higher values for higher virtual fish speeds, consistent with the trend observed in the mean speed (Fig. 3A). Strikingly, while the mean speed of the real fish was already close to the virtual fish’s value, the peak of the PDF aligns almost exactly with it. For all virtual speeds, the real fish has a clear preference for adopting the virtual companion’s speed. Moreover, the real fish maintains consistently the same distance to the wall as the virtual fish (PDF peak at *r*_w_ ≈ 5 cm in Fig. S1B), orients its swimming in the same direction, parallel to the wall (sharp peak of the PDF at *θ*_w_ ≈ 90° in Fig. S1E), and is most frequently at a slightly smaller depth (PDF peaks at *z* ≈ 4 cm in Fig. S1G). The small peak at *θ*_w_ ≈ − 90° corresponds to instances where the real fish swims in the opposite direction to that of the virtual one. Note that trajectories have been normalized so that the virtual fish rotation is anticlockwise (*θ*_w_ *>* 0). The PDF of *z* explains the average depth difference: the real fish tends to explore deeper regions, up to the bowl bottom, particularly when the speed of the virtual fish is higher (long PDF tail up to *z* ≈ 12 cm in Fig. S1G). For all measurements over all virtual speeds, simulations faithfully reproduce the experimental observations.

Figure 4 shows the measures characterizing collective behaviour, and reveals key nuances that modify the idea that the real fish is simply copying the virtual one. First, the PDF of the distance between fish shows a prominent peak around *d* ≈ 2–4 cm for all virtual fish speeds (Fig. 4A), quite smaller than the mean value of 10 cm found for all speeds. Indeed, the PDF also exhibits a long tail going up to *d* ≈ 30 cm, indicating frequent separations where the real fish visits the opposite side of the bowl. This reveals a more complex interaction than simply copying at *d* ≈ 10 cm, as the mean value would have suggested.

**Figure 4:**
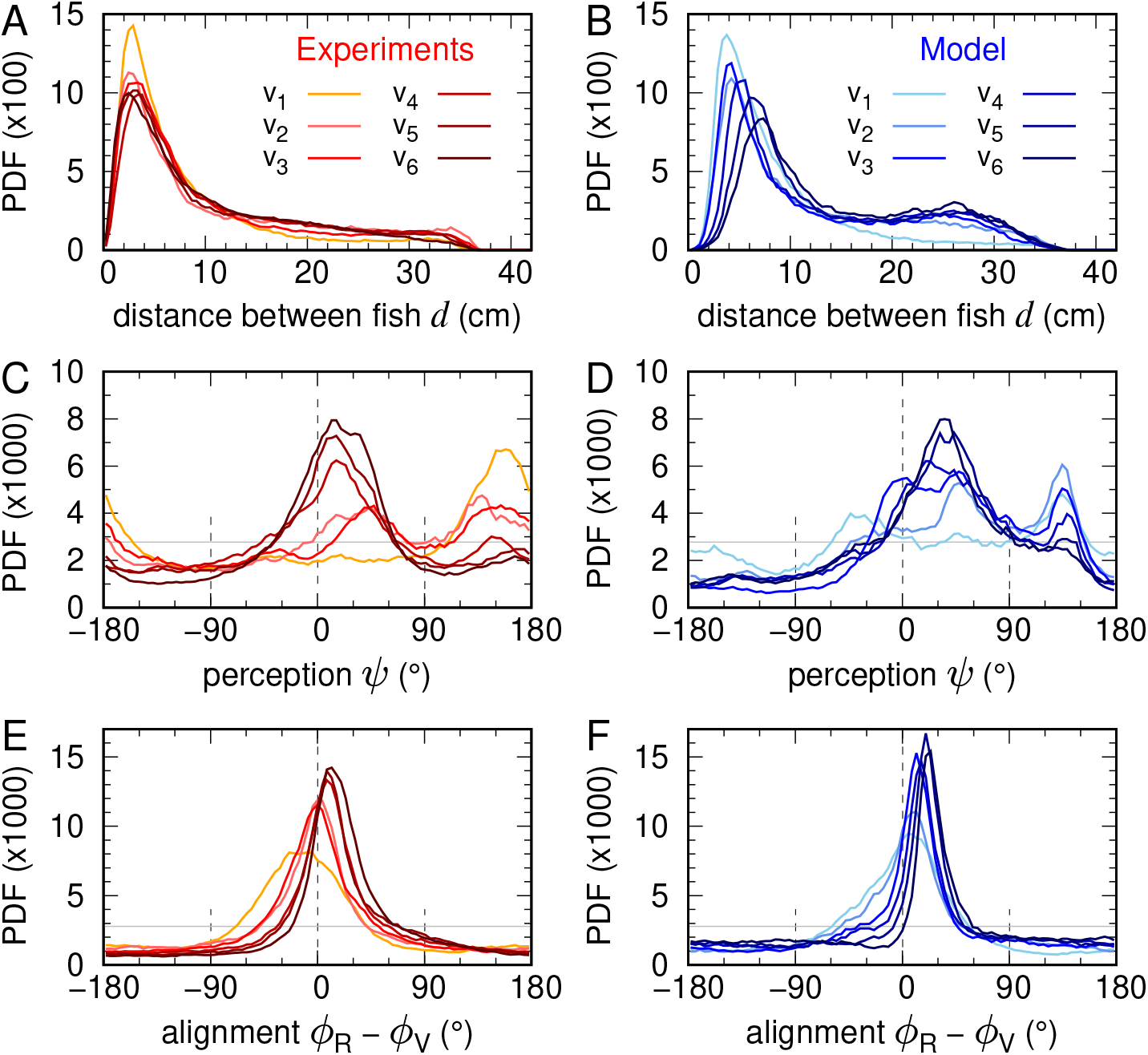
Distributions of pairwise interaction variables across virtual-fish speeds. Probability density functions (PDFs) of (A, B) the distance between fish *d*, (C, D) the angle of perception *ψ*, and (E, F) the alignment between fish (heading-angle difference *ϕ*_R_ − *ϕ*_V_), for the six swimming speeds of the virtual fish (v_1_ to v_6_: 5, 7.5, 10, 11.67, 13.33, 15 cm/s). Experimental results are shown in red and model simulations in blue. In each panel, the six curves correspond to the six virtual-fish speeds, with colour intensity increasing with the virtual fish’s speed.

Another significant indication of complex interaction beyond simple copying is revealed by the PDF of the viewing angle *ψ* of the real fish (Fig. 4C; where the wall is set on the right of the virtual fish). When the virtual fish swims at or below the speed of comfort, the real fish predominantly positions itself ahead of the virtual one (|*ψ*| *>* 90°), whereas at higher speeds, the real fish occupies a follower position (|*ψ*| *<* 60°). Specifically, at the slowest speed (v_1_ = 5 cm/s), the PDF peaks at *ψ* ≈ 160°, meaning that the virtual fish is almost directly behind the real one. For v_2_ = 7.5 cm/s and v_3_ = 10 cm/s, a second peak appears around *ψ* ≈ 45° (ahead and to the side), while at higher speeds, the peak is centred at *ψ* ≈ 20°, indicating that the real fish is clearly behind the virtual one. Thus, at the highest speed (v_6_ = 15 cm/s), the real fish follows the virtual one from the outer side of the virtual fish’s circular trajectory, as illustrated in Fig. 2F, resulting in larger values of *r*_w_ for the real fish.

Finally, the alignment between the real and virtual fish (Fig. 4E) is less pronounced at lower speeds (especially for v = v_1_). Indeed, at the lowest speeds, the real fish frequently overshoots the virtual fish before turning back.

### 3.2 Real fish adjust interaction strengths to virtual conspecific speed

We now use the model to explain the collective behaviours of real—virtual fish pairs for each swimming speed of the virtual fish. For each speed condition (v_1_ to v_6_, the strength of the social interactions and the effects of the bowl’s wall are adjusted to reproduce the behaviour of the real fish. This adjustment was deliberately minimal and consisted solely of applying a specific multiplicative factor to each of the four interaction functions: *f*_Att_, 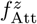, *f*_Ali_ and *f*_w_.

The results of the model simulations for each virtual-fish speed (v_1_ to v_6_) are shown in Fig. 3, Fig. S1 in the electronic supplementary material, and Fig. 4, alongside the corresponding experimental results. In all conditions, the simulations closely capture the observed behavioural patterns, including the fine-scale structure of the PDF of the angle of perception, with a distinct peak near *ψ* ≈ 150° for low virtual fish speeds (Fig. 4D) and the frequent large separations between fish observed across all speeds (Fig. 4B).

Figure 5 summarises how the four multiplicative coefficients vary with the speed of the virtual fish. Both the alignment coefficient and the wall-effect coefficient increase with virtual-fish speed, whereas the strengths of horizontal and vertical attraction decrease. The impact of these coefficients on the shape of the resulting social interaction and wall effect functions is illustrated in the electronic supplementary material (Figure S2). As the speed of the virtual fish adopts larger values, the attraction (Fig. S2A in the plane; Fig. S2D along the *z* axis) and alignment (Fig. S2B) interactions exhibit a smooth but opposite variation in both magnitude and spatial range: attraction is weaker and alignment is stronger for higher speeds of the virtual fish. Wall repulsion (Fig. S2C) also changes smoothly and is stronger for higher speeds of the virtual fish.

**Figure 5:**
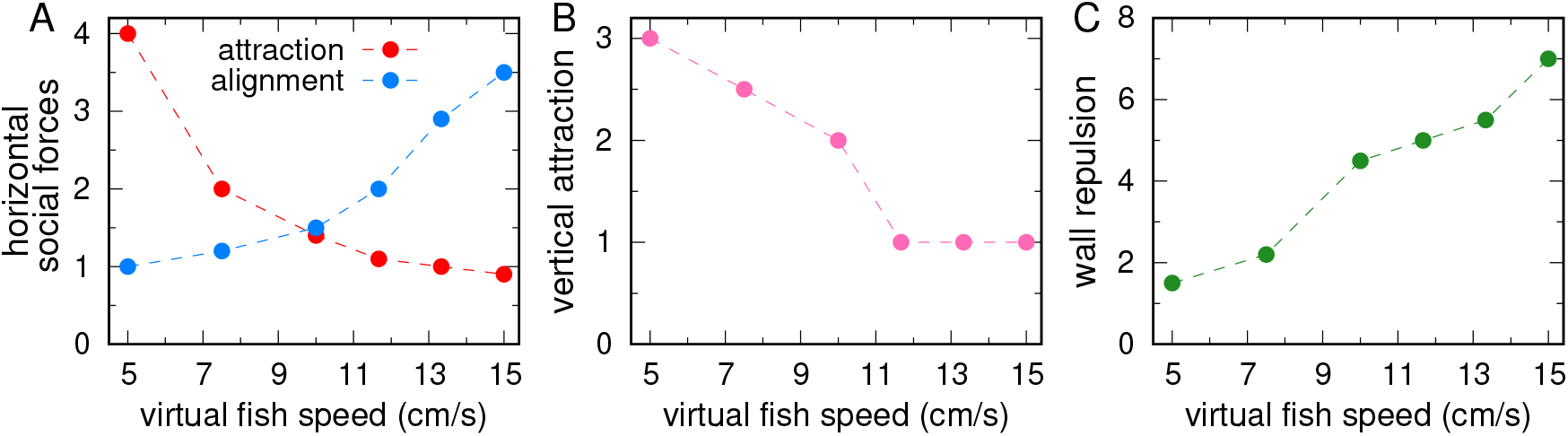
Speed-dependent modulation of interaction forces. Multiplicative coefficients applied to the interaction forces in the model to reproduce the behaviour of the real fish in each of the six speed conditions of the virtual fish (v_1_ to v_6_: 5, 7.5, 10, 11.67, 13.33, 15 cm/s). (A) Coefficients of the horizontal social forces: attraction (red) and alignment (blue), (B) vertical-attraction coefficient, (C) wall-interaction coefficient.

## 4 Discussion

This study examined how a real fish adjusts its behaviour when interacting with a virtual conspecific whose swimming speed is precisely controlled. The central aim was to determine how swimming speed modulates key social interactions, namely attraction, alignment, vertical attraction, and wall avoidance. We quantified these effects using an immersive open-loop virtual-reality system where a real fish follows a virtual partner moving along predefined circular trajectories [18]. We then used a data-driven 3D self-propelled particle model to estimate how the strengths of each interaction vary with speed. The model builds on earlier work that showed that this biohybrid framework captures the primary behavioural forces underlying coordinated swimming [21], allowing us to adjust only multiplicative coefficients to reveal how interaction strengths depend on partner speed. This approach provided a direct and quantitative way to uncover how real fish tune their social interactions when following a faster or slower neighbour.

A first finding is that real fish reduce their attraction to the virtual fish as its swimming speed increases. Both the magnitude and spatial range of attraction decline at higher speeds, as revealed by the fitted multiplicative coefficients and the reconstructed interaction functions. Vertical attraction also decreases with speed, mirroring the trend observed for horizontal attraction. Real fish often swam deeper than the virtual fish at high speed. This behaviour may reduce visual overlap or help stabilise body orientation when turning on curved paths. The behavioural data are consistent with these findings and show that real fish spend more time at larger distances from the virtual fish when it swims rapidly. Reduced attraction at high speed may reflect biomechanical constraints, as close-range following requires rapid turning and frequent acceleration, limitations also reported in studies quantifying speed-dependent turning capacity and inertial constraints in fish and a robotic dummy fish [14]. Similar patterns have been described in natural fish pairs and in groups where higher speed relaxes cohesion by increasing inter-individual spacing [27], reinforcing the view that attraction is context-dependent rather than fixed. The present results therefore demonstrate that fish relax positional coupling at high speed, possibly to preserve stability and minimise energetic or mechanical costs associated with sharp manoeuvres, an interpretation consistent with biomechanical and sensory-motor analyses documenting reduced perceptual accuracy during fast bouts and burst phases [28].

In contrast, alignment becomes stronger as the virtual fish swims faster. The alignment coefficient rises smoothly with speed and the peak of the alignment function increases while extending over a larger spatial range. Stronger alignment at high speed likely facilitates rapid adjustments in heading, reducing the risk of trajectory divergence. Behavioural distributions confirm that fish adopt follower positions more frequently at higher speeds, a spatial configuration that favours alignment-based control. Empirical work shows that increases in speed commonly enhance polarization and shift neighbour positioning from lateral to frontal configurations, thereby strengthening alignment in many fish species [3, 27, 29]. Other studies similarly report speed-selective alignment or selective attention to faster neighbours, mechanisms that can produce speed-induced leadership and asymmetric influence within pairs and groups [15]. Together, these results indicate that fish transition from attraction-dominated interactions at low speed to alignment-dominated interactions at high speed, enabling coherent following when movement dynamics become more demanding.

Swimming speed also influences the magnitude of the wall repulsion, which becomes stronger as speed increases. This trend likely reflects the need to prevent collisions when manoeuvrability decreases at higher speeds, and may serve to maintain safety margins in confined spaces. Together, these trends highlight the joint influence of biomechanical limits and environmental constraints on how fish regulate their position relative to both partner and wall.

The combined results reveal a coherent pattern of speed-dependent tuning of social interactions. Attraction decreases, alignment increases, vertical attraction weakens, and wall repulsion grows with partner speed. These coordinated adjustments illustrate that fish do not rely on static rules, but instead modulate interaction strengths in response to the mechanical and perceptual demands imposed by different movement regimes. These conclusions are fully compatible with theoretical predictions and experimental studies showing that speed is a fundamental control parameter shaping collective states, polarization, and information transfer in fish groups [3, 16, 27, 30]. The biohybrid methodology used here was crucial for revealing these effects, as it ensured controlled variation of a single parameter, the partner speed, while preserving ecologically relevant sensory cues. The modelling framework then reduced the behavioural complexity to a few interpretable coefficients, providing a concise quantitative description of how interaction rules adapt to changing kinematic contexts. This integrated approach mirrors recent efforts to combine behavioural reconstruction, VR-based causal testing, and hybrid real-virtual interactions to identify the functional structure of social rules in fish and other animals [21, 28, 31]. Recent immersive virtual-reality experiments further indicate that coordinated swimming in fish emerges from the joint contribution of speed-dependent spatial interactions and reciprocal temporal coupling, whereby bidirectional feedback between individuals enhances responsiveness to partner manoeuvres and stabilises collective motion [28].

The study focused exclusively on dyads, and larger groups may exhibit additional emergent responses to speed variation. For instance, group-level elongation, polarization, and leadership patterns have been shown to depend sensitively on the distribution of individual speeds and their variability in groups of three or more individuals [14, 27]. The virtual fish followed a simple circular trajectory, which does not capture the diversity of movements seen in natural interactions [21]. Furthermore, adjusting a single coefficient per interaction type provides a parsimonious description but may overlook finer state-dependent effects such as selective attention, reciprocal temporal coupling, or leader-follower transitions that emerge dynamically in real pairs [15, 28]. Despite these constraints, the combination of immersive virtual reality and a data-driven model offers a powerful tool for dissecting how specific variables shape social behaviour. Future work incorporating several virtual fish interacting with one real individual will help determine how general these speed-dependent adjustments are and how, in combination with selective attention, they contribute to collective coordination in natural schools. Extensions of this biohybrid methodology could also clarify whether the speed-dependent tuning observed here scales to the formation of collective states, abrupt changes in polarization, or information propagation observed in larger groups.

## Supporting information

Electronic Supplementary Material

## Ethical Statement

All experimental protocols were approved by the Animal Experimentation Ethics Committee C2EA-01 of the Toulouse Biology Research Federation and were carried out in an approved fish facility (A3155501) under permit APAFIS#27303-2020090219529069 v8 in agreement with the French legislation. All procedures were designed to minimize stress and handling. No animals were sacrificed during this study.

## Data Accessibility

The code and the datasets generated during and/or analysed during the current study are available at: 10.6084/m9.figshare.30584603.

## Acknowledgements

This work was funded by Agence Nationale de la Recherche (ANR-20-CE45-0006-1). RE was partially supported by the Spanish AEI grant PID2020-115088RB-I00. The funders had no role in study design, data collection and analysis, decision to publish, or preparation of the manuscript.

## References

[1] James E. Herbert-Read, Andrea Perna, Richard P. Mann, Timothy M. Schaerf, David J. T. Sumpter, and Ashley J. W. Ward. Inferring the rules of interaction of shoaling fish. Proceedings of the National Academy of Sciences, 108(46):18726–18731, 2011.

[2] Yael Katz, Kolbjørn Tunstrøm, Christos C. Ioannou, Cristián Huepe, and Iain D. Couzin. Inferring the structure and dynamics of interactions in schooling fish. Proceedings of the National Academy of Sciences, 108(46):18720–18725, 2011.

[3] J. Gautrais, F. Ginelli, R. Fournier, S. Blanco, M. Soria, H. Chaté, and G. Theraulaz. Deciphering interactions in moving animal groups. PLOS Computational Biology, 8:e1002678, 2012.

[4] T. Vicsek and A. Zafeiris. Collective motion. Physics Reports, 517(3-4):71–140, 2012. doi: 10.1016/j.physrep.2012.03.004.

[5] D. S. Calovi, A. Litchinko, V. Lecheval, U. Lopez, A. Pérez-Escudero, H. Chaté, C. Sire, and G. Theraulaz. Disentangling and modeling interactions in fish with burst and coast swimming reveal distinct alignment and attraction behaviors. PLOS Computational Biology, 14:e1005933, 2018.

[6] K. Tunstrøm, Y. Katz, C. C. Ioannou, C. Huepe, M. J. Lutz, and I. D. Couzin. Collective states, multistability and transitional behavior in schooling fish. PLoS Computational Biology, 9(2):e1002915, 2013. doi: 10.1371/journal.pcbi.1002915.

[7] J. E. Herbert-Read, S. Krause, L. J. Morrell, T. M. Schaerf, J. Krause, and A. J. W. Ward. The role of individuality in collective group movement. Proceedings of the National Academy of Sciences, 110:13452–13456, 2013. doi: 10.1073/pnas.1309702110.

[8] J. W. Jolles, N. Weimar, T. Landgraf, P. Romanczuk, J. Krause, and D. Bierbach. Group-level patterns emerge from individual speed as revealed by an extremely social robotic fish. Biology Letters, 16:20200436, 2020.

[9] J. W. Jolles, N. J. Boogert, V. H. Sridhar, I. D. Couzin, and A. Manica. Consistent individual differences drive collective behavior and group functioning of schooling fish. Current Biology, 27:2862–2868, 2017. doi: 10.1016/j.cub.2017.08.004.

[10] S. B. Rosenthal, C. R. Twomey, A. T. Hartnett, H. S. Wu, and I. D. Couzin. Revealing the hidden networks of interaction in mobile animal groups allows prediction of complex behavioral contagion. Proceedings of the National Academy of Sciences, 112(15):4690–4695, 2015. doi: 10.1073/pnas.1420068112.

[11] P. Domenici and R. W. Blake. The kinematics and performance of fish fast-start swimming. Journal of Experimental Biology, 200(8):1165–1178, 1997.

[12] S. Marras, S. S. Killen, G. Claireaux, P. Domenici, and D. J. McKenzie. Behavioural and kinematic components of the fast-start escape response in fish: individual variation and temporal repeatability. Journal of Experimental Biology, 214(18):3102–3110, 2011. doi: 10.1242/jeb.056556.

[13] J. E. Herbert-Read, A. Perna, R. P. Mann, T. M. Schaerf, D. J. T. Sumpter, and A. J. W. Ward. Inferring the rules of interaction of shoaling fish. Proceedings of the National Academy of Sciences, 108 (46):18726–18731, 2011. doi: 10.1073/pnas.1109355108.

[14] P. P. Klamser, L. Gómez-Nava, T. Landgraf, J. W. Jolles, D. Bierbach, and P. Romanczuk. Impact of variable speed on collective movement of animal groups. Frontiers in Physics, 9:715996, 2021.

[15] A. Puy, E. Gimeno, J. Torrents, P. Bartashevich, M. C. Miguel, R. Pastor-Satorras, and P. Romanczuk. Selective social interactions and speed-induced leadership in schooling fish. Proceedings of the National Academy of Sciences, 121:e2309733121, 2024.

[16] D. S. Calovi, U. Lopez, S. Ngo, C. Sire, H. Chaté, and G. Theraulaz. Swarming, schooling, milling: Phase diagram of a data-driven fish school model. New Journal of Physics, 16:015026, 2014.

[17] C. Buhl, D. J. T. Sumpter, I. D. Couzin, J. J. Hale, E. Despland, E. R. Miller, and S. J. Simpson. From disorder to order in marching locusts. Science, 312(5778):1402–1406, 2006. doi: 10.1126/science.1125142.

[18] Stéphane Sanchez, Ramón Escobedo, Renaud Bastien, Boris Lenseigne, Audrey Denis, Mathieu Moreau, Maud Combe, Andrew D. Straw, Clément Sire, and Guy Theraulaz. A low-cost immersive virtual reality system to study social interactions and collective behavior in fish. bioRxiv, 2025. doi: 10.1101/2025.05.20.655100. URL https://doi.org/10.1101/2025.05.20.655100. Preprint.

[19] A. Cavagna, I. Giardina, T. Mora, and A. M. Walczak. Physical constraints in biological collective behaviour. Current Opinion in Systems Biology, 9:49–54, 2018. doi: 10.1016/j.coisb.2018.02.006.

[20] Nicholas T. Ouellette. A physics perspective on collective animal behavior. Physical Biology, 19:021004, 2022. doi: 10.1088/1478-3975/ac4805.

[21] R. Escobedo, J. Reynaud, R. Bastien, S. Sanchez, C. Sire, and G. Theraulaz. Closed-loop real-virtual interactions validate 3d model of social coordination in fish. bioRxiv, 2025. doi: 10.64898/2025.12.07.692829. preprint.

[22] Daniel S. Calovi, Alexandra Litchinko, Valentin Lecheval, Ugo Lopez, Alfonso Pérez Escudero, Hugues Chaté, Clément Sire, and Guy Theraulaz. Disentangling and modeling interactions in fish with burst and coast swimming reveal distinct alignment and attraction behaviors. PLoS Computational Biology, 14(4):e1005933, 2018.

[23] Lei Lei, Ramon Escobedo, Clément Sire, and Guy Theraulaz. Computational and robotic modeling reveal parsimonious combinations of interactions between individuals in schooling fish. PLoS Computational Biology, 16(3):e1007194, 2020.

[24] Tingting Xue, Xu Li, Guozheng Lin, Ramón Escobedo, Zhangang Han, Xiaosong Chen, Clément Sire, and Guy Theraulaz. Tuning social interactions’ strength drives collective response to light intensity in schooling fish. PLoS Computational Biology, 19(11):e1011636, 2023.

[25] Weijia Wang, Ramón Escobedo, Stéphane Sanchez, Clément Sire, Zhangang Han, and Guy Theraulaz. The impact of individual perceptual and cognitive factors on collective states in a data-driven fish school model. PLoS Computational Biology, 18(3):e1009437, 2022.

[26] R Escobedo, V Lecheval, V Papaspyros, F Bonnet, F Mondada, Clément Sire, and Guy Theraulaz. A data-driven method for reconstructing and modelling social interactions in moving animal groups. Philosophical Transactions of the Royal Society B, 375(1807):20190380, 2020.

[27] M. I. A. Kent, R. Lukeman, J. T. Lizier, and A. J. W. Ward. Speed-mediated properties of schooling. Royal Society Open Science, 6:181482, 2019.

[28] G. Amichay, L. Li, M. Nagy, and I. D. Couzin. Revealing the mechanism and function underlying pairwise temporal coupling in collective motion. Nature Communications, 15:4356, 2024.

[29] U. Lopez, J. Gautrais, I. D. Couzin, and G. Theraulaz. From behavioural analyses to models of collective motion in fish schools. Interface Focus, 2:693–707, 2012.

[30] S. Mishra, K. Tunström, I. D. Couzin, and C. Huepe. Collective dynamics of self-propelled particles with variable speed. Physical Review E, 86:011901, 2012.

[31] L. Li, M. Nagy, G. Amichay, W. Wang, O. Deussen, D. Rus, and I. D. Couzin. Reverse engineering the control law for schooling in zebrafish using virtual reality. Science Robotics, 10(101), April 2025. doi: 10.1126/scirobotics.adq6784. URL https://doi.org/10.1126/scirobotics.adq6784.

