## Supplementary material for "Swimming speed of schooling fish controls social interaction strength in open-loop immersive virtual reality": Electronic Supplementary Material

### Electronic Supplementary Material for

**This PDF file includes:**

- Supplementary text
- Figures S1, S2
- Tables S1, S2
- Movies S1, S2

### Supplementary text

#### Preprocessing of experimental trajectories

The raw trajectories obtained from experiments were preprocessed to ensure data accuracy and to remove periods of fish inactivity. To preserve temporal continuity, gaps resulting from this filtering process were filled via linear interpolation, provided the missing data spanned less than one second. Gaps exceeding this duration were not reconstructed, resulting in naturally discontinuous trajectories. Moreover, time intervals where a fish remained stationary for 4 seconds or more were removed, as well as segments where the instantaneous speed was higher than 25 cm/s or where the fish maintained a straight, constant-speed swimming for over one second (due to interpolation of missing data in short intervals). These criteria, established from statistical analysis of spontaneous swimming behaviour, effectively removed inactivity phases and tracking artifacts such as abrupt positional jumps. The dataset was further cleaned by excluding the frames in which the reconstructed position fell outside the bowl.

The final trajectories thus consist of multiple consecutive segments. Each segment was then smoothed using a Gaussian kernel convolution to mitigate tracking noise while preserving key kinematic characteristics,

$$x(n\Delta t) = \frac{1}{S_n}\sum_{i=-n}^{n} x\big((n+i)\Delta t\big)K_i, \quad \text{where } K_i = \exp\left[-\left(\frac{i\Delta t}{h}\right)^2\right], \quad S_n = \sum_{i=-n}^{n} K_i, \tag{S1}$$

and $n = 4h/\Delta t$, ensuring negligible kernel weight at $i = n$. The kernel width was set to $h = 0.5\,\text{s}$, achieving effective high-frequency noise reduction with minimal smoothing.

#### Detailed formulation of the model and the interaction functions

The kinematic decomposition of the equations of motion allows us to incorporate the fundamental symmetry constraints that the functional form of the angular terms must verify to consistently reproduce the directional reactions of fish. These constraints preserve the right-left symmetry, which was shown to consistently hold for the considered fish species.

For instance, the rotational force $F_{\text{rot}}$ must include a factor $\sin(\theta_{\text{w}})$ ensuring that a fish approaching the wall with $\theta_{\text{w}} > 0$ (wall to its right) is pushed to turn left with the same intensity as it is pushed to turn right when it approaches the wall with $\theta_{\text{w}} < 0$ (wall to its left). Similarly, the alignment force $F_{\text{Ali}}$ must include a factor $\sin(\Delta\phi)$ favouring a left turn when the neighbour is misaligned to the left ($\Delta\phi > 0$) by increasing the perpendicular acceleration, while if the neighbour is misaligned to the right ($\Delta\phi < 0$), the contribution must be negative to induce a right turn.

These constraints preserve the right-left symmetry, which was shown to consistently hold for the considered fish species.

We assume that, once decomposed along the parallel and perpendicular components of the acceleration, the forces can be separated as products of single-variable functions, so the forces accounting for the effect of the wall are written as,

$$F_{\mathrm{w}}(r_{\mathrm{w}}, \theta_{\mathrm{w}}) = f_{\mathrm{w}}(r_{\mathrm{w}})\, g_{\mathrm{w}}(\theta_{\mathrm{w}}), \tag{S2}$$

$$F_{\mathrm{rot}}(r_{\mathrm{w}}, \theta_{\mathrm{w}}) = f_{\mathrm{rot}}(r_{\mathrm{w}})\, g_{\mathrm{rot}}(\theta_{\mathrm{w}}) \sin(\theta_{\mathrm{w}}), \tag{S3}$$

while those representing social interactions are:

$$F_{\mathrm{Att}}(d, \psi, \Delta\phi) = f_{\mathrm{Att}}(d)\, g_{\mathrm{Att}}(\psi)\, h_{\mathrm{Att}}(\Delta\phi), \tag{S4}$$

$$F_{\mathrm{Ali}}(d, \psi, \Delta\phi) = f_{\mathrm{Ali}}(d)\, g_{\mathrm{Ali}}(\psi)\, h_{\mathrm{Ali}}(\Delta\phi) \sin(\Delta\phi), \tag{S5}$$

$$F^{z}_{\mathrm{Att}}(d, d_z, \psi, \Delta\phi) = f^{z}_{\mathrm{Att}}(d)\, k^{z}_{\mathrm{Att}}(d_z)\, g^{z}_{\mathrm{Att}}(\psi)\, h^{z}_{\mathrm{Att}}(\Delta\phi), \tag{S6}$$

where functions denoted by $g$ and $h$ are even.

Moreover, some of the forces introduced above have a general structure that follows from basic physical considerations. For instance, the magnitude (denoted with lowercase $f$) of the friction should recall the speed to a value $\mathrm{v}_0$ with a relaxation time $\tau_0$ and therefore should have a functional form such as $f_{\mathrm{water}}(\mathrm{v}) = (\mathrm{v}-\mathrm{v}_0)/\tau_0$. Similarly, the force of adaptation should push the fish to match the speed of its neighbour(s), and therefore should be expressed as $f_{\mathrm{adapt}}(\mathrm{v} - \bar{\mathrm{v}}) = (\mathrm{v} - \bar{\mathrm{v}})/\tau_{\mathrm{adapt}}$. Regarding vertical speed, its mean in a bounded bowl should be zero, so the vertical friction should have the form $f^{z}_{\mathrm{water}}(\mathrm{v}_z) = \mathrm{v}_z/\tau_z$.

This results in the following stochastic differential system,

$$\begin{aligned} a_{\|} = {} & -f_{\mathrm{water}}(\mathrm{v}) - f_{\mathrm{adapt}}(\mathrm{v} - \bar{\mathrm{v}}) - f_{\mathrm{w}}(r_{\mathrm{w}})\, g_{\mathrm{w}}(\theta_{\mathrm{w}}) \cos\theta_{\mathrm{w}} \\ & + f_{\mathrm{Att}}(d)\, g_{\mathrm{Att}}(\psi)\, h_{\mathrm{Att}}(\Delta\phi) \cos\psi + \eta_{\|}, \end{aligned} \tag{S7}$$

$$\begin{aligned} a_{\perp} = {} & f_{\mathrm{w}}(r_{\mathrm{w}})\, g_{\mathrm{w}}(\theta_{\mathrm{w}}) \sin\theta_{\mathrm{w}} + f_{\mathrm{rot}}(r_{\mathrm{w}})\, g_{\mathrm{rot}}(\theta_{\mathrm{w}}) \sin\theta_{\mathrm{w}} + f_{\mathrm{Att}}(d)\, g_{\mathrm{Att}}(\psi)\, h_{\mathrm{Att}}(\Delta\phi) \sin\psi \\ & + f_{\mathrm{Ali}}(d)\, g_{\mathrm{Ali}}(\psi)\, h_{\mathrm{Ali}}(\Delta\phi) \sin\Delta\phi + \eta_{\perp}, \end{aligned} \tag{S8}$$

$$a_z = -f^{z}_{\mathrm{water}}(\mathrm{v}_z) - f^{z}_{\mathrm{comfort}}(z) + f^{z}_{\mathrm{Att}}(d)\, k^{z}_{\mathrm{Att}}(d^z)\, g^{z}_{\mathrm{Att}}(\psi)\, h^{z}_{\mathrm{Att}}(\Delta\phi) + \eta_z, \tag{S9}$$

where $\bar{\mathrm{v}}$ is the mean speed of the neighbour(s) of the focal fish.

This system is solved numerically with the Euler–Maruyama scheme and appropriate initial conditions based on experimental data.

The interaction functions, assumed to be the product of single-variable interaction functions, can be empirically reconstructed from the experimental trajectories by expressing each one as a combination of polynomial expansions and Fourier series. The parameters of these expressions are then optimized with an iterative process that minimizes the error between the experimentally measured acceleration of a real fish and the acceleration predicted by the model.

This procedure is applied to extract the analytical form of the single-variable functions from the experimental sessions conducted with two real fish. They are presented in the next section.

### Analytical expressions of the extracted interaction functions

$$f_{\mathrm{water}}(\mathrm{v}) = C_{\mathrm{v}}^{(1)} + C_{\mathrm{v}}^{(2)}(\mathrm{v}-\mathrm{v}_0) + C_{\mathrm{v}}^{(3)}(\mathrm{v}-\mathrm{v}_0)^2 + C_{\mathrm{v}}^{(4)}(\mathrm{v}-\mathrm{v}_0)^3, \tag{S10}$$

$$f_{\mathrm{adapt}}(\mathrm{v}) = C_{\mathrm{v}}^{(5)}\mathrm{v} + C_{\mathrm{v}}^{(6)}\mathrm{v}^2 + C_{\mathrm{v}}^{(7)}\mathrm{v}^3, \tag{S11}$$

$$f_{\mathrm{w}}(r_{\mathrm{w}}) = C_{f_{\mathrm{w}}}^{(1)} \exp\left[-\frac{r_{\mathrm{w}}}{C_{f_{\mathrm{w}}}^{(2)}} - \left(\frac{r_{\mathrm{w}}}{C_{f_{\mathrm{w}}}^{(3)}}\right)^2\right] - C_{f_{\mathrm{w}}}^{(4)}, \tag{S12}$$

$$f_{\mathrm{rot}}(r_{\mathrm{w}}) = C_{f_{\mathrm{rot}}}^{(1)} \exp\left[-\frac{r_{\mathrm{w}}}{C_{f_{\mathrm{rot}}}^{(2)}} - \left(\frac{r_{\mathrm{w}}}{C_{f_{\mathrm{rot}}}^{(3)}}\right)^2\right] - C_{f_{\mathrm{rot}}}^{(4)}, \tag{S13}$$

$$g_{\mathrm{w}}(\theta_{\mathrm{w}}) = 1 + \sum_{m=1}^{6} C_{\mathrm{w}}^{(m)} \cos(m\theta_{\mathrm{w}}), \quad g_{\mathrm{rot}}(\theta_{\mathrm{w}}) = 1 + \sum_{m=1}^{6} C_{\mathrm{rot}}^{(m)} \cos(m\theta_{\mathrm{w}}), \tag{S14}$$

$$f_{\mathrm{Att}}(d) = \frac{d - C_{f_{\mathrm{Att}}}^{(3)}}{C_{f_{\mathrm{Att}}}^{(1)}} \times \left[1 + \left(\frac{d}{C_{f_{\mathrm{Att}}}^{(2)}}\right)^2\right]^{-C_{f_{\mathrm{Att}}}^{(4)}}, \tag{S15}$$

$$\text{If } d < d_{\mathrm{c}}^{\mathrm{Att}},\ f_{\mathrm{Att}}(d) = f_{\mathrm{Att}}(d) - \left[\left(\frac{d_{\mathrm{c}}^{\mathrm{Att}}}{d}\right)^2 - 1\right] \frac{d_{\mathrm{c}}^{\mathrm{Att}}}{C_{f_{\mathrm{Att}}}^{(1)}}; \quad \text{If } f_{\mathrm{Att}} < f_{\mathrm{Att}}^{\mathrm{min}},\ f_{\mathrm{Att}} = f_{\mathrm{Att}}^{\mathrm{min}}, \tag{S16}$$

$$f_{\mathrm{Ali}}(d) = \frac{d - C_{f_{\mathrm{Ali}}}^{(3)}}{C_{f_{\mathrm{Ali}}}^{(1)}} \times \left[1 + \left(\frac{d}{C_{f_{\mathrm{Ali}}}^{(2)}}\right)^2\right]^{-C_{f_{\mathrm{Ali}}}^{(4)}}, \tag{S17}$$

$$E_{\mathrm{Att}}(\psi) = 1 + \sum_{m=1}^{6} C_{\mathrm{Att}}^{(m)} \cos(m\psi), \quad E_{\mathrm{Ali}}(\psi) = 1 + \sum_{m=1}^{6} C_{\mathrm{Ali}}^{(m)} \cos(m\psi), \tag{S18}$$

$$G_{\mathrm{Att}}(\Delta\phi) = 1 + \sum_{m=1}^{6} D_{\mathrm{Att}}^{(m)} \cos(m\Delta\phi), \quad G_{\mathrm{Ali}}(\Delta\phi) = 1 + \sum_{m=1}^{6} D_{\mathrm{Ali}}^{(m)} \cos(m\Delta\phi), \tag{S19}$$

$$f_{\mathrm{water}}^{z}(\mathrm{v}_z) = C_{\mathrm{v}_z}^{(1)}\mathrm{v}_z + C_{\mathrm{v}_z}^{(2)}\mathrm{v}_z^2 + C_{\mathrm{v}_z}^{(3)}\mathrm{v}_z^3 \tag{S20}$$

$$f_z(z) = C_{f_z}^{(1)} + C_{f_z}^{(2)}(z - z_0) + C_{f_z}^{(3)}(z - z_0)^2 + C_{f_z}^{(4)}(z - z_0)^3 + C_{f_z}^{(5)}(z - z_0)^4, \tag{S21}$$

$$f_{\mathrm{Att}}^{z}(d_z) = C_{f_{\mathrm{Att},z}}^{(1)} d_z + C_{f_{\mathrm{Att},z}}^{(2)} d_z^2 + C_{f_{\mathrm{Att},z}}^{(3)} d_z^3 + C_{f_{\mathrm{Att},z}}^{(4)} d_z^4, \tag{S22}$$

$$f_{\mathrm{Att,z,d}}(d) = C_{f_{\mathrm{Att,z,d}}}^{(1)} \left[1 + \left(\frac{d}{C_{f_{\mathrm{Att,z,d}}}^{(2)}}\right)^{C_{f_{\mathrm{Att,z,d}}}^{(3)}}\right]^{-1}, \tag{S23}$$

$$E_{\mathrm{Att}}^{z}(\psi) = 1 + \sum_{m=1}^{6} C_{\mathrm{Att,z}}^{(m)} \cos(m\psi), \quad G_{\mathrm{Att}}^{z}(\Delta\phi) = 1 + \sum_{m=1}^{6} D_{\mathrm{Att,z}}^{(m)} \cos(m\Delta\phi). \tag{S24}$$

### Supplementary Figures

Figure S1 presents the PDFs of individual behavioural variables of the real fish, comparing the experimental results (left column, in red) to the model simulations (right column, in blue).

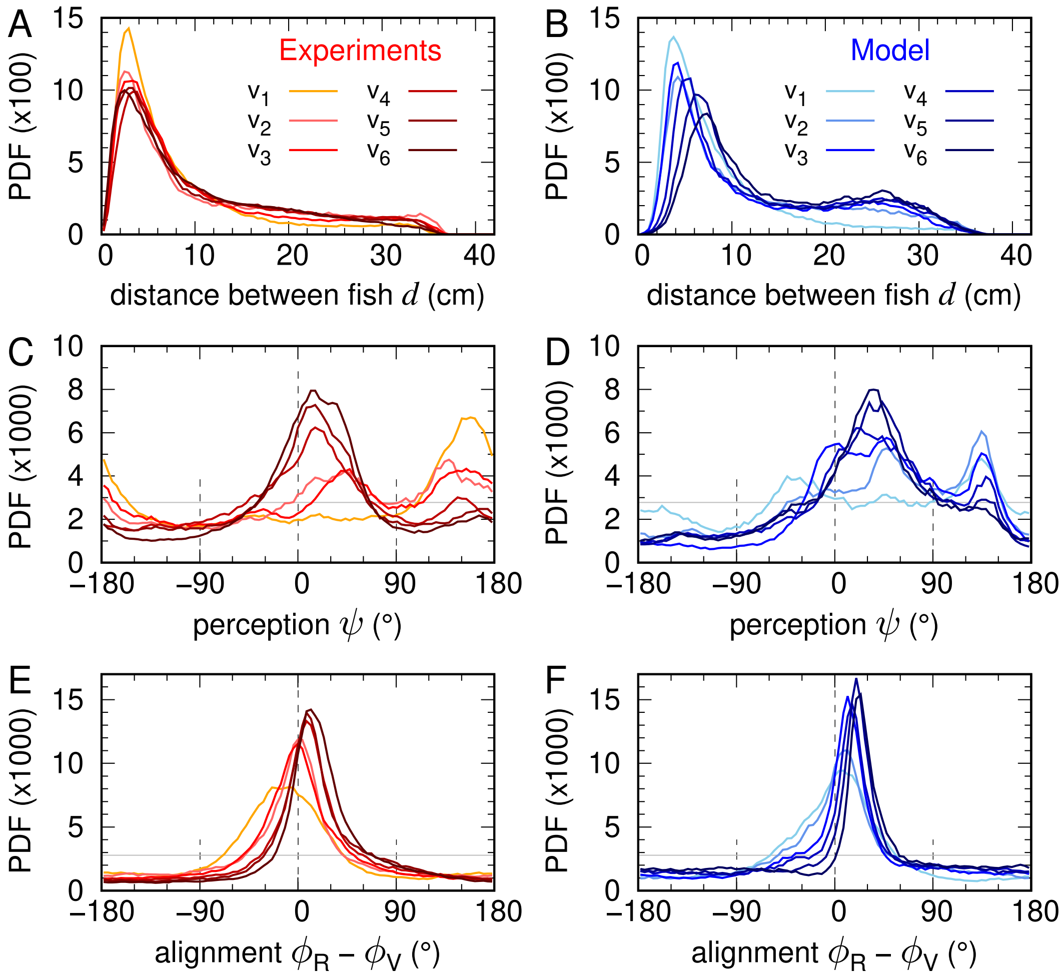


Figure S1: **Distributions of individual behavioural metrics across virtual-fish speeds.** Probability density functions (PDFs) of key behavioural variables of the real fish for the six swimming speeds of the virtual fish ($v_1$ to $v_6$: 5, 7.5, 10, 11.67, 13.33 and 15 cm/s). Experimental results are shown in red and model simulations in blue. (A, B) Swimming speed $v$, (C, D) distance to the wall $r_w$, (E, F) angle of incidence to the wall $\theta_w$, and (G, H) swimming depth $z$. In each panel, colour intensity increases with the virtual fish's speed.

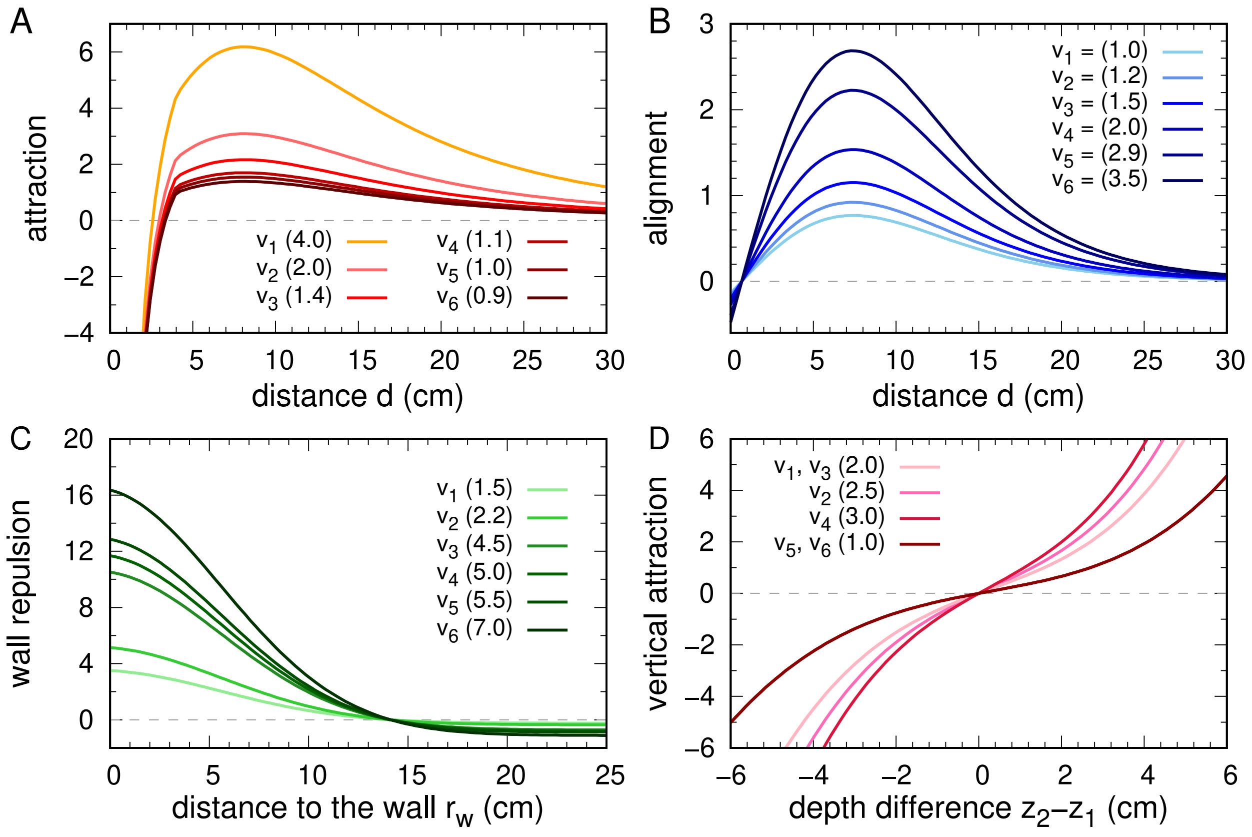


Figure S2: **Interaction functions across virtual-fish speed conditions**. Social interaction functions used in the model simulations for each swimming speed of the virtual fish ($v_1$ to $v_6$: 5, 7.5, 10, 11.67, 13.33 and 15 cm/s). (A) Attraction and (B) alignment as functions of the horizontal inter-individual distance $d$, (C) wall repulsion as a function of the horizontal distance to the border $r_\mathrm{w}$, and (D) vertical attraction as a function of the vertical separation between fish $z_2 - z_1$. In each panel, the six curves correspond to the six virtual-fish speeds, with colour intensity increasing with the virtual fish's speed. Values in parentheses indicate the multiplicative coefficients applied in each condition.

### Supplementary Tables

Table S1: Linear regression analysis, experiments vs model

| | Experiments | | Model | | | Experiments | | Model | |
|---|---|---|---|---|---|---|---|---|---|
| | **$R^2$** | **p-value** | **$R^2$** | **p-value** | | **$R^2$** | **p-value** | **$R^2$** | **p-value** |
| v | 0.941 | 0.0003 | 0.946 | 0.0002 | $z$ | 0.877 | 0.016 | 0.811 | 0.17 |
| $r_w$ | 0.109 | 0.59 | 0 | 0.97 | $d$ | 0.049 | 0.36 | 0.424 | 0.007 |

Table S2: Model parameters

| **Parameter** | **Symbol** | **Value** |
|---|---|---|
| Parallel acceleration noise | $\sigma_{\parallel}$ | 3.4 |
| Perpendicular acceleration noise | $\sigma_{\perp}$ | 2.9 |
| Vertical acceleration noise | $\sigma_z$ | 2.9 |
| Noise correlation time | $\tau$ | 0.15 |
| Typical speed, friction coefficient | $v_0$ | 12 |
| Multiplicative coefficient of: | | |
| speed adaptation | $Coef\_Adapt$ | 0 |
| friction | $Coef\_Friction$ | 0.25 |
| vertical friction | $Coef\_Friction_z$ | 0.75 |
| depth preference | $Coef\_f_z$ | 0.5 |
| attraction | $Coef\_f_{att}$ | Fig. 7A |
| alignment | $Coef\_f_{ali}$ | Fig. 7B |
| wall interaction | $Coef\_f_w$ | Fig. 7C |
| vertical attraction | $Coef\_f_{att,z}$ | Fig. 7D |

### Supplementary Movies

#### S1 Movie. Behavioural response of a real fish to the circular motion of a virtual conspecific with a low swimming speed.

Video excerpts of an experiment with a real fish interacting with the anamorphic projection of a virtual conspecific in the experimental bowl of radius 250 mm. The virtual fish moves along a uniform circular trajectory with a constant speed of 5 cm/s, at a constant depth of 5 cm below the water surface, and at a constant distance of 5.4 cm to the edge of the tank. Top-Left: user interface allowing the real-time visualization of the trajectories of the real and virtual fish (in the $xy$- and $xz$-planes) and the instantaneous modification of the parameters of the model driving the virtual fish. Bottom-Left: real-time 3D tracking of the real fish. Right panel: anamorphic rendering of the virtual fish projected onto the acrylic bowl by the rendering application according to the 3D position of the real fish.

#### S1 Movie. Behavioural response of a real fish to the circular motion of a virtual conspecific with a high swimming speed.

Video excerpts of an experiment with a real fish interacting with the anamorphic projection of a virtual conspecific in the experimental bowl of radius 250 mm. The virtual fish moves along a uniform circular trajectory with a constant speed of 15 cm/s, at a constant depth of 5 cm below the water surface, and at a constant distance of 5.4 cm to the edge of the tank. Top-Left: user interface allowing the real-time visualization of the trajectories of the real and virtual fish (in the $xy$- and $xz$-planes) and the instantaneous modification of the parameters of the model driving the virtual fish. Bottom-Left: real-time 3D tracking of the real fish. Right panel: anamorphic rendering of the virtual fish projected onto the acrylic bowl by the rendering application according to the 3D position of the real fish.
